# An SNF2 helicase-like protein links mitotic transcription termination to sister chromatid resolution

**DOI:** 10.1101/2022.11.21.517340

**Authors:** Catarina Carmo, João Coelho, Rui Silva, Alexandra Tavares, Ana Boavida, Paola Gaetani, Rui Gonçalo Martinho, Raquel A. Oliveira

## Abstract

Mitotic chromatin is largely assumed incompatible with transcription due to changes in the transcription machinery and chromosome architecture. However, the mechanisms of mitotic transcriptional inactivation and their interplay with chromosome assembly remain largely unknown. By monitoring ongoing transcription in *Drosophila* early embryos, we reveal that eviction of nascent mRNAs from mitotic chromatin occurs after substantial chromosome compaction and is not promoted by condensin I. Instead, we show that the timely removal of transcripts from mitotic chromatin is driven by the SNF2 helicase-like protein Lodestar (Lds), identified here as a modulator of sister chromatid cohesion defects. In addition to transcriptional termination, we uncovered that Lds cooperates with Topoisomerase 2 to ensure efficient sister chromatid resolution and mitotic fidelity. We conclude that mitotic transcriptional termination is not a passive consequence of cell cycle progression and/or chromosome compaction but occurs via dedicated mechanisms with functional parallelisms to sister chromatid resolution.

## Introduction

Mitotic chromosome assembly is essential for faithful genome distribution during mitosis. Defects in this process can damage dividing chromosomes, causing DNA breaks and genetic instability. This process encompasses changes in the compaction level and the topology of genome organisation, with concomitant global shutdown of most transcriptional activity (Gottesfeld & Forbes, 1997; Piskadlo & Oliveira, 2016). Nevertheless, how the structural changes that occur on mitotic chromatin impact its transcriptional state, while long postulated (Gottesfeld & Forbes, 1997), remains an open question.

The architectural reshaping of mitotic chromosomes is primarily mediated by the condensin complexes. Condensins are ring-like protein complexes known to extrude DNA molecules in an ATP-dependent manner (Davidson *et al*, 2019; Ganji *et al*, 2018; Kim *et al*, 2019), thereby leading to chromosome compaction. This process occurs with the concurrent resolution of sister chromatids, driven by two primary mechanisms: 1) the removal of protein-mediated linkages around DNA molecules, by the cohesin destabilising factor WAPL, which removes the molecular glue from chromosome arms (Gandhi *et al*, 2006; Gimenez-Abian *et al*, 2004; Kueng *et al*, 2006; Losada *et al*, 1998; Sumara *et al*, 2000); 2) the resolution of topological DNA-DNA links (e.g. catenations), by Topoisomerase 2 (Top2), which introduces a dsDNA break in one of the chromatids, allows for strand passage, and reseals the break (Pommier *et al*, 2016). Condensin-mediated chromosome compaction and removal of cohesin from chromosome arms aid in this process by positioning DNA molecules in a manner that biases Top2 activity towards the decatenation of DNA-DNA intertwines (Baxter *et al*, 2011; Piskadlo & Oliveira, 2017; Piskadlo *et al*, 2017; Sen *et al*, 2016).

Alongside these structural changes, mitotic chromosomes switch off most of their transcriptional activity (Gottesfeld & Forbes, 1997). In some cell types, such as the rapid divisions of early *Drosophila* embryos, entry into mitosis dictates abortion of ongoing transcription (Rothe *et al*, 1992; Shermoen & O’Farrell, 1991). The mechanisms that drive transcription termination/abortion and how they relate to the structural changes remain poorly understood. Recent studies propose that cohesin release contributes to the removal of active RNA Polymerase II (PolII) from chromosomes since abnormal cohesin retention is sufficient to accumulate active PolII along chromosome arms (Perea-Resa *et al*, 2020). Chromosome compaction is also assumed to actively or passively contribute to the switch off of transcription (Gottesfeld & Forbes, 1997), although this remains untested. The loop extrusion activity of SMC complexes (Davidson *et al*., 2019; Ganji *et al*., 2018; Kim *et al*.,2019) (in particular of condensin) is a likely candidate to stall PolII or drive its eviction. Accordingly, condensin is often found at transcription termination sites in yeast mitotic chromosomes and is proposed to contribute to transcript release (Nakazawa *et al*, 2019). Moreover, mitotic bookmarking by TATA-binding protein (TBP) depends on local condensin inactivation (Xing *et al*, 2008). However, studies in bacteria showed that condensins can bypass the transcriptional machinery despite a reduction of speed upon their encounter (Brandao *et al*, 2019). Thus, it remains unclear if condensin loading on mitotic chromosomes influences transcription termination in metazoans.

In parallel to changes in chromosome organisation, regulation of the transcriptional machinery offers a direct layer of regulation of transcriptional switch-off. For example, *in vitro* studies support that biochemical inactivation of the basal transcription machinery, driven by Cdk1-mediated phosphorylation, prevents *de novo* transcription initiation (Akoulitchev & Reinberg, 1998; Bellier *et al*, 1997; Gebara *et al*, 1997; Long *et al*, 1998; Segil *et al*, 1996) but the prevalence of this inhibition *in vivo* remains unclear. Studies in human cells proposed that the transcription termination Factor 2 (TTF2) drives the removal of active PolII in mitosis (Liu *et al*, 1998) whereas the transcription initiation inhibitor Gdown1 was shown to gain access to chromatin solely during mitosis to repress mitotic transcription (Ball *et al*, 2022). Recent studies have also proposed that specific mechanisms ensure promoter clearance upon mitotic entry, mediated by PolII-mediated activation of Top1, to release topological stress (Wiegard *et al*, 2021) or hyperactivation of P-TEFb, to enhance transcriptional elongation (Liang *et al*, 2015).

However, and despite these findings, it remains unclear how these pathways impact transcription dynamics relative to cell cycle progression and concomitant changes in chromosome architecture, as this process has never been monitored in real-time. Using livecell imaging of ongoing transcription in dividing *Drosophila* early embryos, we show that condensin-I mediated compaction does not drive mitotic transcription inactivation (MTI). Instead, we reveal that the SNF2 helicase-like protein Lds, the *Drosophila* orthologue of human TTF2 (Liu *et al*., 1998), ensures prompt eviction of nascent RNAs from mitotic chromatin. In parallel to MTI, Lds also aids DNA decatenation of sister chromatid intertwines. We conclude Lds holds a dual function in mitotic chromosomes, uncovering unforeseen functional parallelisms between mitotic transcription inactivation and sister chromatid resolution.

## Results

### A genetic screen for modifiers of cohesion loss uncovers *lodestar* as a strong suppressor of cohesion defects

We have previously established a genetic screen to find modulators of sister chromatid cohesion (Silva *et al*, 2018). This screen uses the morphology defects in *Drosophila* wings obtained upon depletion of the cohesin stabiliser separation anxiety (san) (Hou *et al*, 2007; Ribeiro *et al*, 2016; Rong *et al*, 2016; Williams *et al*, 2003). Changes in the wing morphology phenotypes are then used to unbiasedly identify conditions that could either enhance or suppress the defects associated with cohesion loss (Silva *et al*., 2018). In a subsequent search for additional cohesion modifiers, using the same approach, we uncovered the SNF-2-like helicase lodestar (lds), the putative orthologue of the human TTF2 (Liu *et al*., 1998), as a modulator of the defects associated with cohesion loss. When san RNAi was driven by the wing imaginal disc blade region-specific driver Nubbin Gal4 (nub-Gal4), flies ecloded with significant adult wing abnormalities (Fig 1 and (Ribeiro *et al*., 2016; Silva *et al*., 2018)). However, co-expression of san RNAi with lds RNAi (but not with luciferase-RNAi) efficiently suppressed these abnormalities and wings were significantly closer to normal morphology. Hence, depletion of Lds is likely a potent suppressor of cohesion defects observed after san RNAi (Fig 1).

**Figure 1.**
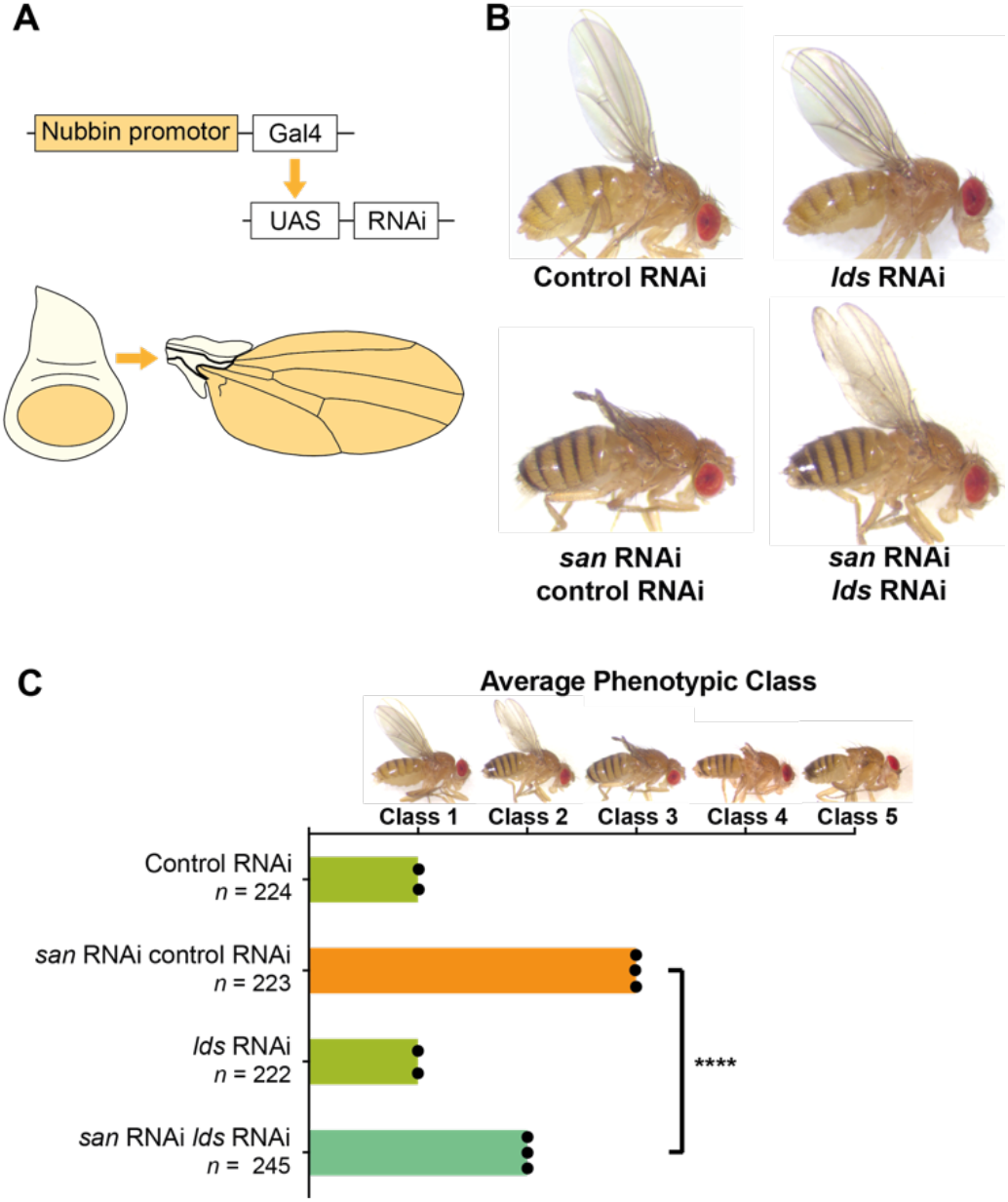
Depletion of Lds strongly suppresses the defects associated to cohesin loss. **(A)** Tissue-specific RNAi in the pouch of the larvae wing imaginal discs using the nubbin-Gal4 driver and the upstream activating sequence (UAS)/Gal4 system. **(B)** Representative images of *Drosophila* adult wings that resulted from larvae wing imaginal discs expressing a control RNAi (mCherry RNAi), a *lds* RNAi, or co-expressing *san* RNAi with control RNAi (san RNAi control RNAi) or with *lds* RNAi (san RNAi lds RNAi). **(C)** Quantification of *Drosophila* adult wing phenotypes expressing individual control RNAi or *lds*RNAi transgenes (light green bars), co-expressing *san* RNAi and control RNAi (orange bar), or coexpressing *san* RNAi and *lds* RNAi (blue-green bar). Classes used for phenotypic quantification were previously described in (Silva *et al*., 2018), and a representative example for each class is shown on top of the graph: class 1 (wild-type wings); class 2 (adult wings with weak developmental defects); class 3 (adult wings with moderate developmental defects; *san* RNAi-like wing phenotype); class 4 (highly abnormal adult wings); and class 5 (absence or vestigial adult wings). Representative images shown in (B) of adult wings co-expressing *san* RNAi and control RNAi correspond to class 3, whereas wings co-expressing *san* RNAi and *lds* RNAi correspond to class 2. Results are the mean of three independent biological replicas for adult wing phenotypes of *Drosophila* co-expressing *san* RNAi with control RNAi or with *lds* RNAi and two independent experiments for wing phenotypes expressing individual RNAi transgenes (control RNAi or *lds* RNAi). The results of independent experiments are represented by black dots. Class scoring average (standard deviation): control RNAi (1.0±0) (n=224), lds RNAi (2.0±0) (n=222), san RNAi control RNAi (2.996±0.006) (n=223), san RNAi lds RNAi (2.0±0) (n=245). ****p < 0.0001; one-way ANOVA with Bonferroni’s multiple comparison test; *n* represents the total number of scored flies.

Lds has been previously described as a maternal-effect gene whose loss of function mutations are associated to mitotic defects during early embryogenesis (Girdham & Glover, 1991). Additionally, a dominant gain of function allele was found to impair meiotic chromosome segregation (Szalontai *et al*, 2009). However, it remains unknown how this protein contributes to the fidelity of nuclear division.

### Lds is associated with mitotic chromatin and required for mitotic fidelity across various tissues

We next sought out to investigate if Lds is also required for the fidelity of mitosis in somatic tissues. We used live-cell imaging analysis to probe for the accuracy of mitosis upon RNAi-mediated depletion of Lds in the developing wing disc. We used nub-Gal4 to drive RNAi for lds (alone), specifically in the wing pouch. We found that in the absence of Lds, the nuclear division is slightly delayed and severely compromised, with a high frequency of anaphase bridges at mitotic exit (Fig 2A-C). Our results confirm that Lds is required for mitotic fidelity in developing wing epithelia, similar to what was reported for embryonic syncytial divisions ((Girdham & Glover, 1991) and Fig S1).

**Figure 2.**
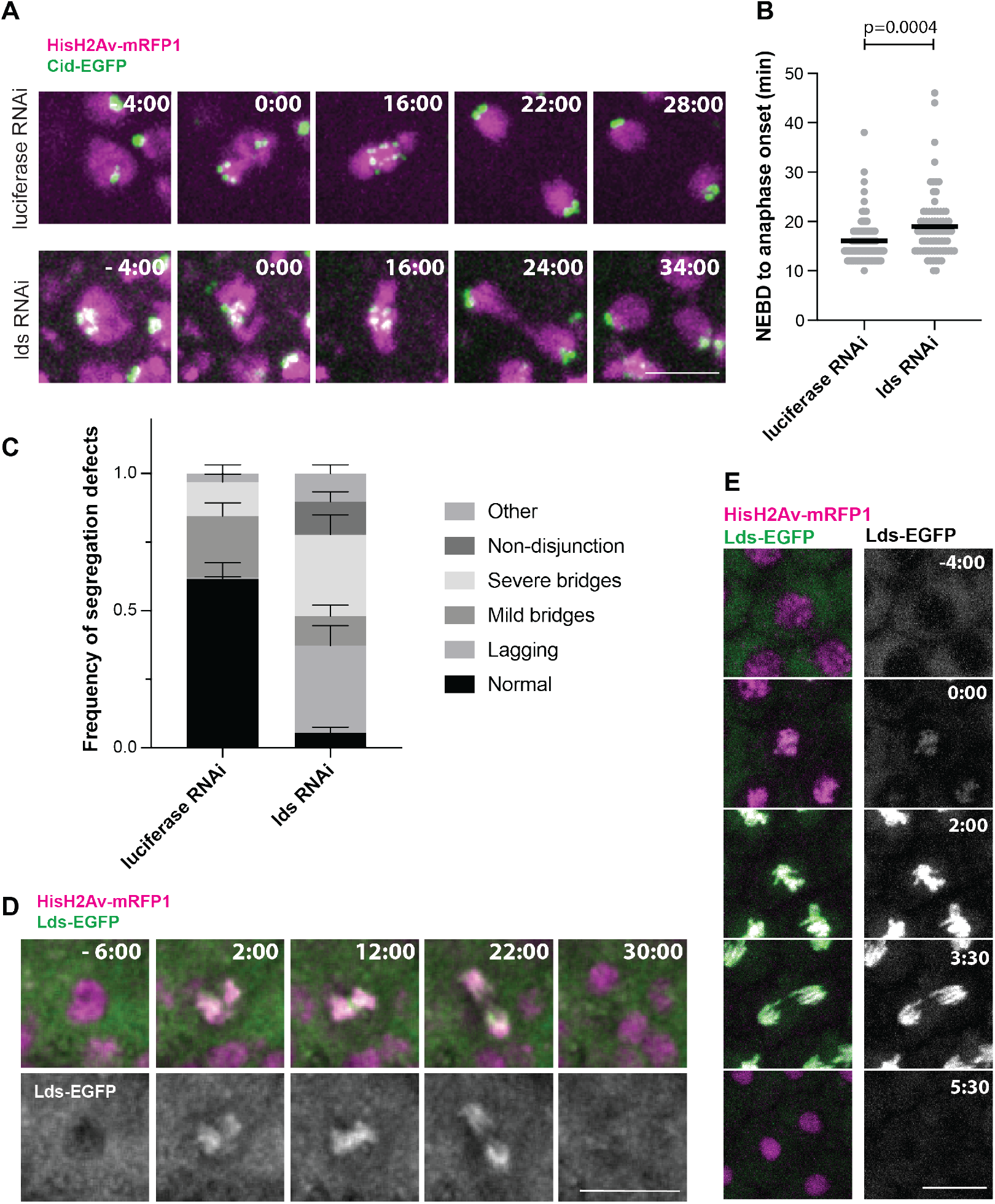
Lds is associated with mitotic chromatin and required for mitotic fidelity. **(A)** Representative images of cells in the wing disc pouch upon nub-Gal4-mediated RNAi for luciferase (top) or lds (bottom). Cells also express HisH2AvD-mRFP1 (magenta) and centromeric marker cid-EGFP (green); Times (min:sec) are relative to NEBD; scale bar is 5 μm and applies to all images. **(B)** Mitotic timing of defined from NEBD to anaphase onset; each dot represents a different cell (n=77 cells from at least 6 independent movies per condition); black lines the mean; statistical analysis performed using the non-parametric Mann-Whitney test. **(C)** Frequency of segregation defects during mitosis in *Drosophila* wing discs. Graph depicts the average frequency of segregation defects observed upon luciferase or Lds depletion, represented as Mean ± SEM (n= 9 discs luciferase RNAi; n= 7 discs lds RNAi; average of 11 cells measured per disc) **(D,E)** Localisation of Lds-EGFP (green) throughout mitosis in *Drosophila* wing discs (**D**) and *Drosophila* syncytial embryos (**E**). Times are relative to NEBD, flies also express HisH2AvDmRFP1 (magenta). Scale bars are 10 μm and apply to all images.

In order to gain further insight into the mitotic functions of Lds, we generated flies carrying a C-terminal EGFP-fusion at the endogenous lds locus (using Crisp-Cas9-based genome editing) and assessed its localisation in *Drosophila* dividing tissues. In both wing discs and syncytial embryos, we observed that Lds is excluded from chromatin during interphase but gains access to mitotic chromatin around the time of nuclear envelope breakdown (NEBD) (Fig 2C,D). The chromatin levels of Lds are maximised during metaphase and decay as cells exit mitosis. These results suggest that Lds acts specifically on mitotic chromatin to ensure faithful genome partition. We, therefore, sought to investigate how Lds contributes to mitotic fidelity.

### Lds mediates the timely release of RNA transcripts from mitotic chromatin

*In vitro* studies have previously demonstrated that Lds (also known as factor 2) has transcription termination activity (Price *et al*, 1987; Xie & Price, 1997; Xie & Price, 1996, 1998). However, this activity has never been demonstrated *in vivo*, and it remains to be determined how it relates to a putative role in chromosome segregation. We focused our analysis on establishing whether transcription termination upon mitotic entry could be defective in the absence of Lds. For this, we used *Drosophila* early embryos as a prime model system. Despite the reduced level of transcription (solely restricted to a minor wave of zygotic transcription (Tadros & Lipshitz, 2009)), several tools allow for efficient monitoring of transcriptional dynamics by live cell imaging. We used a system that enables the visualisation of a reporter transcript that carries MS2 loops at its 5’ and is expressed under the early zygotic gene *hunchback (hb)* promoter (Garcia *et al*, 2013). Upon transcription initiation in flies carrying MCP-EGFP, MCP binds the MS2 loops, and the GFP signal is readily detected at the transcription site, enabling the analysis of transcription in real-time (Garcia *et al*., 2013). Using this approach, we monitored the time of transcript release relative to mitotic progression and chromosome compaction. We find that in wild-type embryos, complete removal of the labelled nascent transcripts from chromatin was only observed following NEBD, considerably after the initiation of chromosome condensation (Fig 3A,B and S3). This finding suggests that transcript release may be mediated by unknown mechanisms that operate shortly after NEBD.

**Figure 3.**
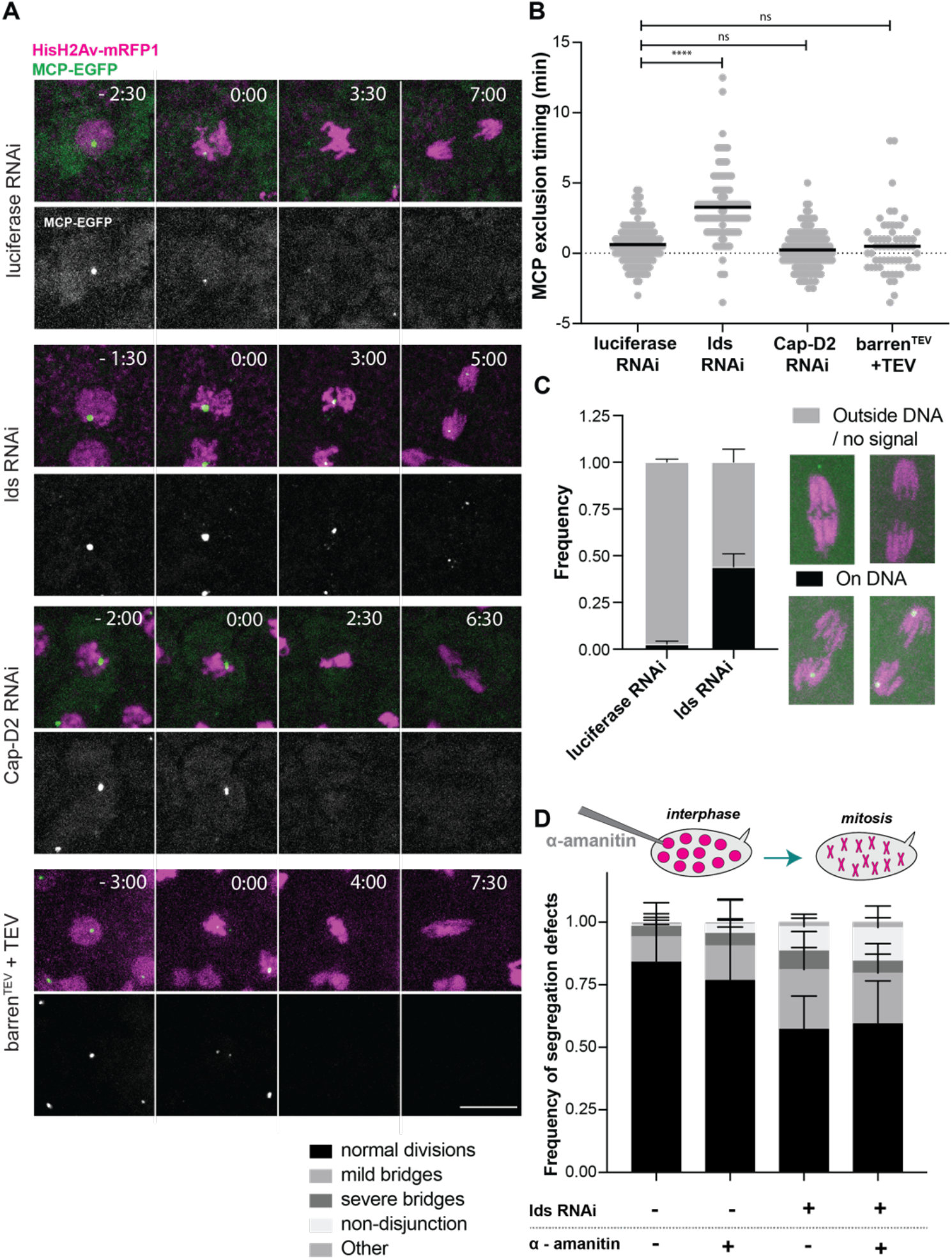
Lds, but not condensin I, is required for timely release of nascent transcripts in mitosis. **(A)** Analysis of ongoing transcription monitored by MCP-EGFP labelled nascent transcripts on a reporter locus containing MS2 loops (green), in the refered experimental conditions. Flies also express H2AvD-mRFP1 (magenta). Scale bar is 10 μm and applies to all images. **(B)** Time of clearance of MCP-labelled transcripts from mitotic chromatin, relative to nuclear envelope breakdown. Each dot represents a single cell and black bars the mean; n= 140 (luciferase RNAi), 122 (Lds RNAi), 150 (Cap-D2 RNAi) and 63 (barren^TEV^ after injection of TEV protease) nuclei, derived from 5-12 individual embryos per experimental condition. **(C)** Frequency of anaphase figures with MCP-EGFP signals (labelling nascent transcripts) observed on/off mitotic chromatin. Graph depicts average from 15 (luciferase) and 17 (lds) independent movies. Error bars are SEM. **(D)** Segregation defects upon inhibition of transcription (microinjection with 1 mg/ml alpha-amanitin in early interphase) in luciferase and lds RNAi. Graph depicts mean ± SEM of the frequencies of defects scored in 11 (luciferase RNAi, H2O and lds RNAi, amanitin) or 8 (lds RNAi, H_2_O and Luc RNAi, amanitin) independent embryos; a total of ~800 nuclei were scored per condition.

Then, we asked if Lds could drive transcript release in these divisions. We monitored the kinetics of nascent mRNAs disappearance from chromatin in embryos depleted of Lds. We used nanos Gal4 (nos-Gal4) driver to induce Lds depletion in the germline, which, similar to what we observed in the wing, leads to severe mitotic defects in these early embryonic divisions (Fig S1), that can be rescued by ectopic addition of Lds (Fig S2A). When looking at the dynamics of ongoing transcription in these embryos, we observed that nascent mRNAs remain attached to mitotic chromatin for much longer in the absence of Lds, relative to controls, and full transcript release could only be detected on average ~3.5 min after NEBD (Fig 3A,B). This delay is partially restored upon ectopic addition of mRNA coding for an RNAi-resistant form of lds (Fig S2B). Notably, while anaphases from control embryos rarely retain nascent mRNAs bound to chromatin, upon lds RNAi, we detected nascent mRNAs still attached in ~44% of anaphases, sometimes remaining throughout the entire cell division cycle (Fig 3C). This localisation is, in most cases, symmetrically placed and consistent with the reporter’s location (i.e. ~1/3 of the left arm of chromosome 3). These observations imply that the detected mRNAs are still present at the active transcript site in the later stages of mitosis. We, therefore, conclude that Lds is critical for the active removal of nascent transcripts during mitosis. This finding supports the working hypothesis that removal of engaged transcripts occurs via dedicated machinery, and not simply as a by-product of the structural changes that occur on the mitotic chromatin.

To confirm this notion, we then asked whether condensin I complex loading, enhanced around the time of NEBD (Oliveira *et al*, 2007), also contributes to transcripts release. We tested this idea by analysing the time of removal of MCP-labelled transcripts from chromatin in embryos depleted for one of the condensin I subunits, Cap-D2. The prediction is that if condensin I activity drives transcripts eviction, its depletion should alone induce a delay in nascent RNA removal. In contrast to this expectation, we observed that RNAi for Cap-D2 did not impose any noticeable delay in nascent mRNA eviction relative to controls (Fig 3). Similar results were obtained upon acute inactivation of the kleisin subunit prior to mitotic entry (TEV-mediated cleavage of the condensin I subunit Barren (Piskadlo *et al*., 2017)). Using this approach, and although the frequency of severe anaphase bridges and mitotic defects was very high (Piskadlo *et al*., 2017), these changes in chromosome organisation were not associated to any delay in the timing of transcripts release (Fig 3). Altogether, these results suggest that loop extrusion mediated by condensin I complexes does not contribute to the eviction of RNA PolII from chromatin during mitosis, and support the hypothesis that timely termination of transcription during mitosis relies on specific mechanisms, driven by the helicase-like protein Lds.

### Transcription termination failures are not the sole cause of the mitotic defects associated to the loss of Lds

Considering the high frequency of mitotic defects observed in embryos depleted of Lds, we asked whether abnormally trapped PolII could be the source for the observed errors. We reasoned that if the segregation defects associated to loss of Lds derive from transcriptionally-dependent events, transcription inhibition should be sufficient to restore mitotic fidelity. To test this notion, we performed transcription inactivation experiments. We microinjected embryos with 1 mg/ml alpha-amanitin in early interphase, which is sufficient to abolish transcription, as evidenced by the absence of MS2-labelled reporter RNAs in the subsequent interphase (Fig S4). Transcription inhibition in wild-type embryos did not compromise mitotic fidelity. This finding implies that, in contrast to prior reports (Liang *et al*, 2008; Staudt *et al*,2006), ongoing transcription is not required for mitotic fidelity, at least within the short time frames of this experimental set-up.

Next, we used the same approach to inhibit transcription in lds RNAi embryos. This analysis revealed that in the presence of transcription (control H2O injection), lds RNAi embryos present ~43 % of abnormal anaphase figures. Upon alpha-amanitin injection, the frequency of these defects is not significantly reduced (41%). These findings imply that the presence of trapped nascent mRNAs on mitotic chromatin is not the primary source of defects that compromise mitotic fidelity in Lds-depleted embryos.

### Lds cooperates with Topoisomerase 2 in sister chromatid resolution

Since transcription inhibition does not rescue the segregation defects observed upon depletion of Lds, we reasoned that those mitotic defects were likely to result from transcription-independent events. We thus hypothesised that Lds might hold additional roles in mitotic fidelity beyond transcriptional termination. We focused on the fact that the most prominent phenotype was a high frequency of chromatin bridges to investigate whether Lds contributes to the resolution of sister chromatids.

To test this, we probed for potential genetic interactions of lds with other conditions known to be involved in the resolution of sister chromatids throughout mitosis. We envisioned two possible ways Lds contributes to the resolution of sister chromatids. One way could be via the stabilisation of cohesin-mediated linkages at the metaphase-anaphase transition, thereby hindering the efficient release of this proteinaceous glue. The second would imply a contribution to the removal of DNA-DNA ties (catenations), catalysed by topoisomerase 2 with the help of condensin I (Piskadlo & Oliveira, 2017). Following this rationale, we employed the same strategy presented above and used the modulation of wing-morphology defects as a readout for potential genetic interactions. Despite the high percentage of mitotic defects, depletion of Lds alone does not cause detectable adult wing-morphology abnormalities, possibly due to compensatory proliferation pathways that ensure tissue homeostasis (Ryoo *et al*, 2004). We next probed for the effect of co-depletion of Lds with other players known to contribute to sister chromatid resolution. We found that removal of Lds could enhance the phenotype associated with overexpression of a modified version of cohesin where the interfaces between Smc3 and Rad21 are covalently linked (Eichinger *et al*, 2013). This covalent fusion promotes cohesin stability by preventing WAPL-mediated release and was previously shown to induce moderate defects in wing morphology (Ribeiro *et al*., 2016) (Fig 4A). The resulting defects were more severe if this fusion is expressed with concomitant depletion of Lds (Fig 4A). Similarly, co-depletion of Lds was also an effective enhancer of the defects associated with the impaired removal of DNA-DNA catenations, including RNAi for the condensin subunit Cap-D2 and RNAi for Top2 (Fig 4A).

**Figure 4.**
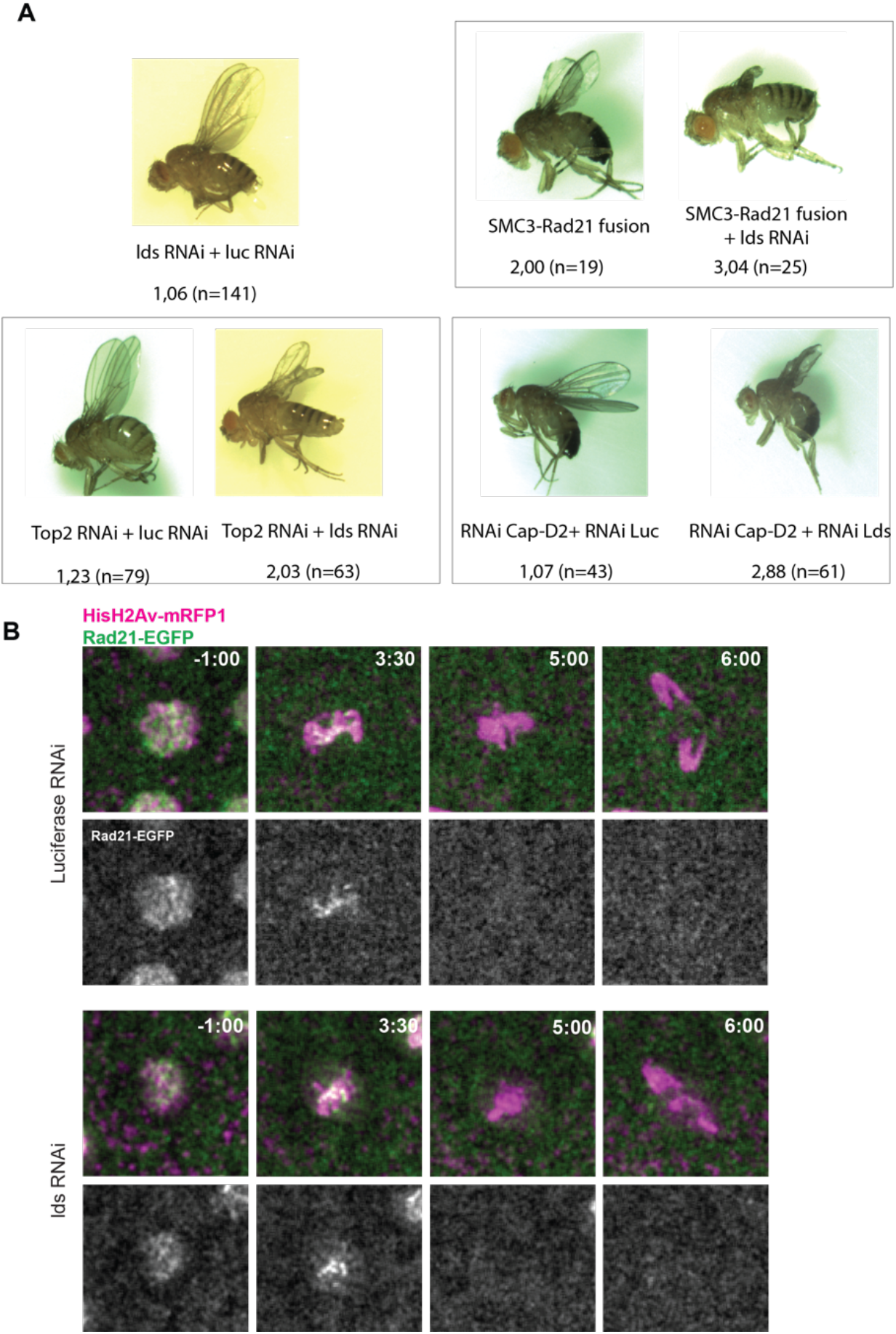
Lds genetically interacts with various players in sister chromatid resolution but does not control cohesin levels in mitosis. **(A)** Wing morphology phenotypes obtained depletion of Lds combined with control co-depletion (luciferase RNAi) or expression/depletion of chromosome architecture modulators. Wing morphology defects were classified according to their severity (1= normal wings and 5= no wing); depicted numbers indicate the average morphology grade and the number of flies counted, from at least 2 independent experiments. **(B)** Live cell imaging analysis of the kinetics of Rad21-EGFP (green) removal from mitotic chromosomes in luciferase RNAi and lds RNAi conditions. DNA is marked with H2AvD-mRFP1 (magenta); times are relative to NEBD; scale bar is 10 μm and applies to all images.

These findings support the hypothesis that Lds is involved in the efficient resolution of sister chromatids. To distinguish whether it works through the release of cohesin and/or DNA catenations, we first monitored the dynamics of the cohesin removal in the presence or absence of Lds. These results revealed that upon depletion of Lds, and similarly to control strains, the cohesin kleisin subunit Rad21-EGFP was mainly detected at the centromeric regions and not along chromosome arms (as observed upon WAPL removal (Oliveira *et al*, 2014)) (Fig 4B). These findings confirm that the removal of cohesin complexes along chromosome arms is unperturbed. Moreover, we could not detect any delay in the kinetics of Rad21-EGFP removal at the metaphase to anaphase transition (Fig 4B), supporting an efficient Separase-mediated cleavage of cohesin. Altogether, these results support that Lds does not enhance or stabilise cohesin complexes on DNA in a way that could potentially preclude the efficient removal of cohesin-mediated linkages.

We thus focused on investigating how Lds could contribute to the removal of DNA catenations, either directly or via interaction with other modulators. We first probed for the presence of Condensin I and Top2 on mitotic chromosomes upon Lds removal. We used strains expressing endogenously-tagged Barren-EGFP (Kleinschnitz *et al*, 2020) and Top2-EGFP (developed here, see Materials and Methods). In both cases, we found that Top2-EGFP and Barren-EGFP were efficiently recruited to mitotic chromosomes (Fig 5 A and B), indicating that Lds is not required for the chromatin targeting of these critical players in chromosome resolution. Interestingly, we observed a significant enrichment of both Top2 and Condensin I subunit Barren at the site of the anaphase bridges observed upon depletion of Lds (Fig 5 A and B). These findings imply that DNA bridges caused by lds knockdown retain the machinery responsible for DNA resolution. Reciprocally, we find that DNA bridges that result from improper sister chromatid resolution induced by other means (e.g. RNAi for the condensin I subunit Cap-D2) retain high levels of Lds, as evidenced by the increased amount of Lds-EGFP at those sites (Fig 5C). These results suggest that the bridges observed upon Lds loss are likely caused by unresolved DNA catenations, resulting in the hyper-recruitment of their resolution machinery at mitotic exit. Interestingly, the presence of Lds in chromatin bridges caused by other means suggests that Lds is by itself part of this resolution mechanism.

**Figure 5.**
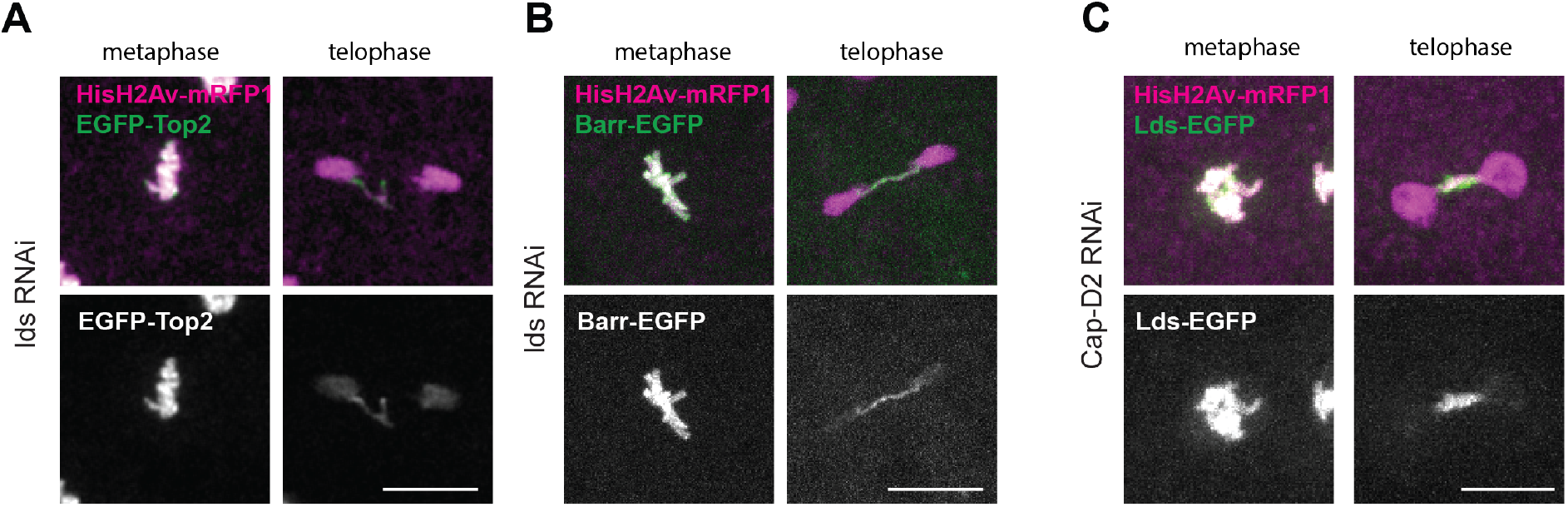
Lds localizes to DNA bridges. (**A, B**) Chromatin localisation of chromosome assembly factors (Top2 (**A**) and Condensin I subunit Barren (**B**)) upon loss of Lds in metaphase cells and anaphase figures where DNA bridges could be identified. DNA is marked with H2AvD-mRFP1 (magenta); scale bars are 10 μm and apply to all images. (**C**) Localisation of Lds-EGFP (green) in embryos depleted of condensin I subunit Cap-D2; DNA is marked with H2AvD-mRFP1 (magenta); scale bars are 10 μm and apply to all images.

To further test a potential cooperation between Lds and the primary enzyme responsible for DNA decatenation, Top2, we sought to probe for a possible interplay between these two proteins on mitotic chromatin. For this, we established conditions to trap Top2 on mitotic chromatin artificially: embryos were microinjected with UbcH10^DN^ to induce metaphase arrest (Oliveira *et al*, 2010) and subsequently microinjected with the Top2 inhibitor ICRF-193. Under these conditions, we observed a sharp increase in the amount of Top2-EGFP detected, despite the overall decompaction of mitotic chromatin (Fig 6B). We then used the same assay to monitor the behaviour of Lds-EGFP upon Top2 inhibition. We found that upon ICRF-193 injection, there was a sharp increase in chromosome-bound Lds-GFP levels (~2.5 fold) (Fig 6C). Notably, the rise in Lds-GFP does not follow the same kinetics as Top2. It increases faster than Top2 itself, reaching a steady-state on equimolar concentrations at later time points. These findings imply that Lds is sensitive to Top2 inhibition and is either co-enriched on mitotic chromosomes via direct binding to Top2 and/or is recognising topological changes on DNA induced upon Top2 inhibition.

**Figure 6.**
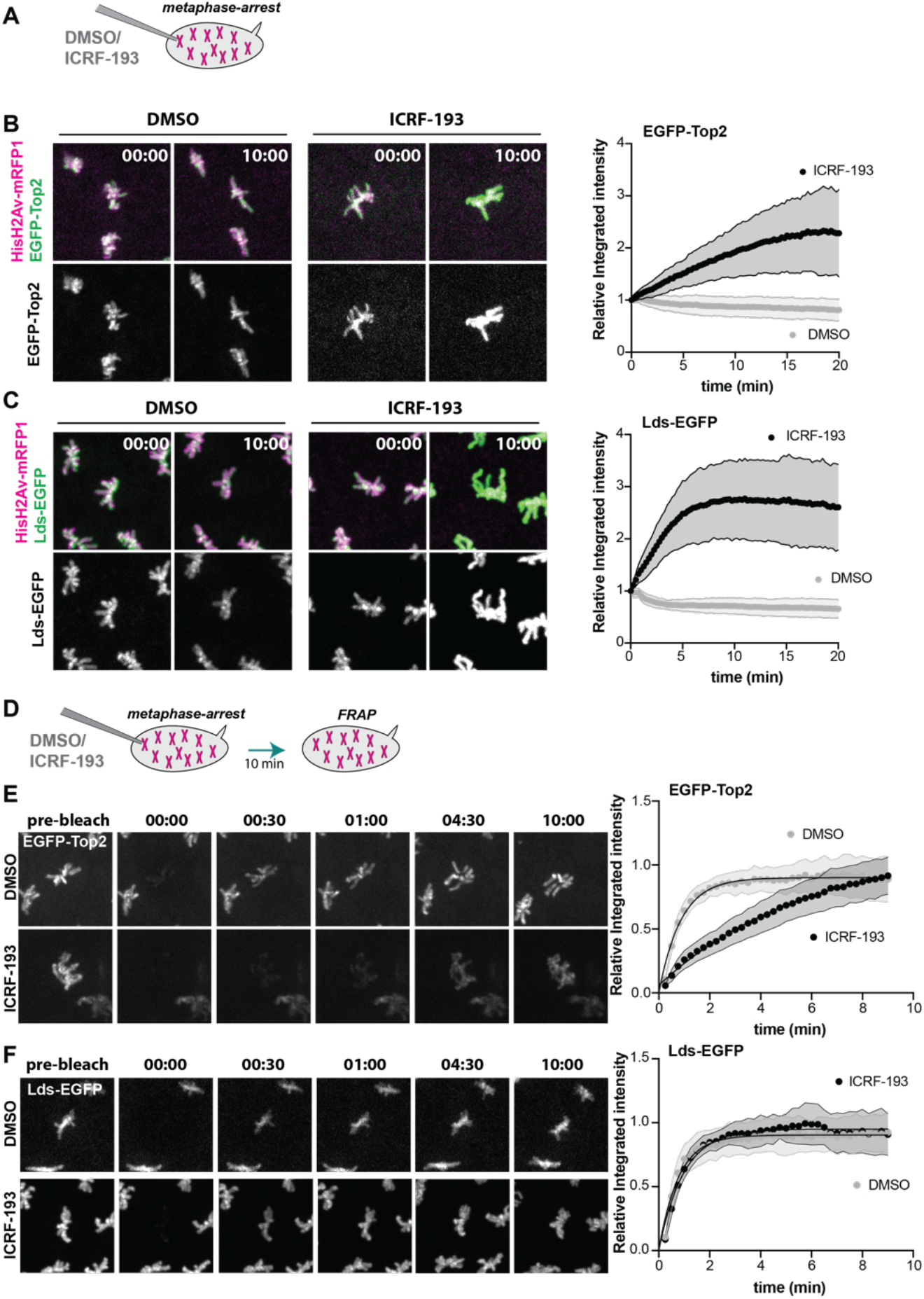
Lds chromatin association is sensitive to Top2 inhibition. **(A-C)** Levels of EGFP-Top (**B**) and Lds-EGFP (**C**) loading on mitotic chromatin of metaphase arrested embryos (UbcH10^C114S^ injection) upon subsequent injection of DMSO/ICRF-193; graphs depict integrated intensities normalized to the first time point, represented as mean (dots) ± SD (grey area); n=54 (Lds+ ICRF-193, Lds+DMSO, Top2 + ICRF-193) or 47 (Top2 + DMSO), derived from 8-9 independent embryos per experimental condition. **(D-F)** FRAP analysis with and without ICRF-193 treatment for EGFP-Top2 (**E**) and Lds-EGFP (**F**). Metaphase-arrested embryos were microinjected with DMSO/ICRF-193 and an entire metaphase plate was bleached 10 min after. Graphs depict recovery of fluorescence signal over time. Dots represent average and grey areas SD; Top2: n= 14 (DMSO) and 18 (ICRF-193) metaphases; Lds: n= 11 (DMSO) and 9 (ICRF-193) metaphases, derived from 3-7 independent embryos per experimental condition.

To distinguish between these two possibilities, we monitored the dynamic behaviour of both proteins using Fluorescence Recovery After Photobleaching (FRAP) with and without ICRF-193 treatment. For this analysis, the fluorescence of an entire metaphase plate was bleached, and the fluorescence recovery was subsequently monitored. This analysis revealed that Top2 displays a highly dynamic association with chromatin (t_1/2_= 36.6 ± 10.8 sec), in accordance with previous reports (Christensen *et al*, 2002; Tavormina *et al*, 2002). A similar analysis on

Lds demonstrates that Lds-EGFP displays a comparable dynamic turnover (t_1/2_= 32.4 ± 5.7 sec). Importantly, we observed that treatment of ICRF-193 increased the half-time of Top2 recovery significantly (t_1/2_= 4.43 ± 1.51 min, p<0.0001 One-way ANOVA), evidencing the differential mode of binding of this molecule to chromatin in the presence of the inhibitor. In contrast, Lds retains its high turnover rate (t_1/2_= 40.14 ± 8.94 sec), indicating it is not co-trapped with Top2 on chromosomes. Altogether, these results suggest that Lds is likely sensitive to changes in the DNA molecule induced by Top2 inhibition, which may aid in the efficient resolution of sister chromatid intertwines.

### Lds has a dual function in mitosis

Considering the dual role observed for Lds on mitotic transcription termination and sister chromatid resolution, we next asked if these two functions are interdependent or represent two distinct actions required during mitosis. To address this, we probed whether transcript release depends on Top2 activity. We monitored the time of MCP-labelled transcripts disappearance from chromatin, in situations of impaired Top2, achieved by microinjection of the Top2 inhibitor ICRF-193 before mitotic entry. This treatment resulted in a significant increase the MCP-EGFP signals in both luciferase and lds RNAi, possibly due to transcription stalling caused by Top2 inhibition/trapping (Fig 7A,-C). However, ICRF-193 injection does not change the timing of transcript release, and a significant delay is solely detected in the absence of Lds, independently of the Top2 activity state (Fig 7A,D). These results imply that Top2 activity is not required for efficient transcriptional termination in these embryos.

**Figure 7.**
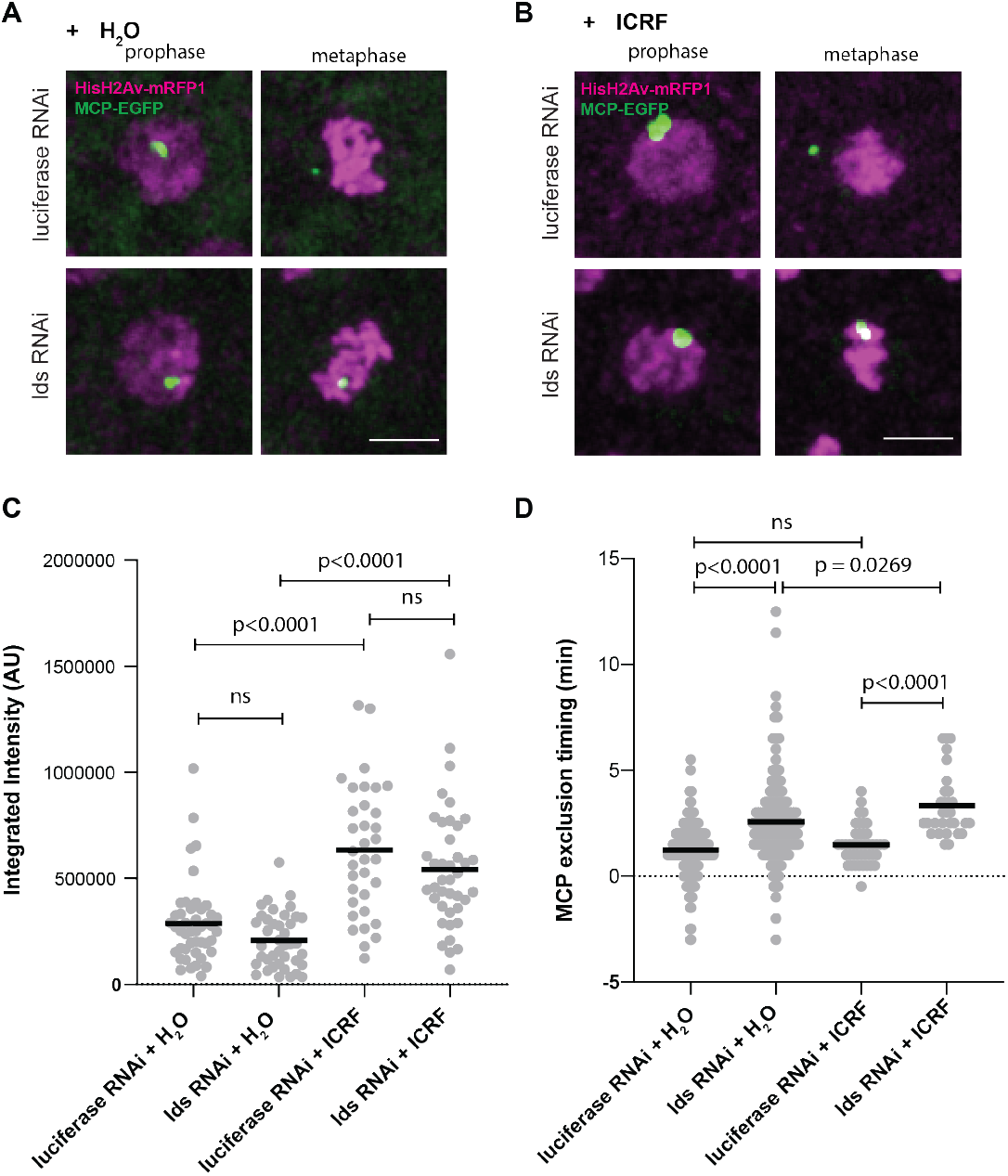
Top2 is dispensable for timely release of mitotic transcripts from chromatin. (**A,B**) Representative images of mitotic nuclei in prophase (1 min before NEBD) and metaphase (1 min before anaphase onset), upon RNAi for luciferase/lds, microinjected with water (**A**) or ICRF-193 (**B**). Scale bars are 5 μm and refer to all images. (**C**) Integrated intensity for the MCP-EGFP signal measured 1 minute before nuclear envelope breakdown with and without ICRF-193 injection. (luciferase RNAi + H_2_O n= 45 nuclei from 7 independent embryos; lds RNAi + H_2_O n= 40 nuclei from 8 embryos; luciferase RNAi + ICRF n= 34 nuclei from 7 embryos; lds RNAi + ICRF-193 n= 40 nuclei from 8 embryos). Statistical analysis was performed using Kruskall-Wallis test, using Dunn’s test for multiple comparisons. (**D**) Time (min) of disappearance of the MCP-EGFP signal from chromatin in the referred conditions, relative to NEBD; n= 196 (luciferase RNAi + H_2_O), 205 (lds RNAi + H2O), 43 (luciferase RNAi + ICRF) and 30 (lds RNAi + ICRF-193) nuclei, derived from at least 3 independent embryos per experimental condition.

## Discussion

Here we uncovered the depletion of lds as a strong suppressor of the defects associated with cohesion loss. We showed that Lds contributes to mitotic fidelity, as a dual-function chromatin factor: it ensures prompt transcription termination and efficient sister chromatid resolution. We further show that removal of mitotic transcripts from chromatin does not rely on Top2 strand passage activity or efficient chromosome decatenation. These findings suggest that Lds may modulate mitotic chromatin to facilitate both processes. Notably, cohesin has also been involved in these two activities, which can explain the suppressor effect on wing morphology. Specifically, the presence of cohesin is known to preclude efficient sister chromatid resolution, possibly by keeping sisters in such proximity that favours sister chromatid intertwining as opposed to their resolution (Farcas *et al*, 2011; Sen *et al*., 2016). Cohesin retention is also required and sufficient for active transcription on mitotic chromosomes (Perea-Resa *et al*., 2020). Hence, our findings uncovered unprecedented links between these two processes that re-shape chromatin during mitosis both at the structural and functional levels.

Classic views have postulated that mitotic transcription inhibition (MTI) would be a passive consequence of mitosis, either due to changes in chromosome organisation and/or by biochemical changes imposed by the rise in cdk1 activity (Gottesfeld & Forbes, 1997). In contrast with these dogmas, our results suggest that MTI is driven by an active mechanism that ensures prompt transcriptional shutdown upon mitotic entry. Note that in the absence of Lds, chromosomes can still condense and progress through mitosis with regular timings (or with slight delays). These findings imply that transcripts can remain engaged with mitotic chromatin despite the chromosomal compaction and high Cdk1 activity. Collectively, these findings support that MTI is a more active and regulated process than previously anticipated.

The need for dedicated machinery to support such mechanisms may have evolved to facilitate mitotic progression. A tempting hypothesis is that pervasive transcription can compromise mitotic fidelity. Indeed, some evidence refers to increased mitotic defects upon chronic perturbation of putative MTI players, although these defects have never been analysed in great detail. For example, RNAi for TTF2 or Gdown1 leads to an increase in the presence of binucleated cells or p53 activation after aberrant mitosis (Ball *et al*., 2022; Jiang *et al*, 2004), whereas impairment of PolII clearance through P-TEFb removal leads to delays in the progression of cell division (Liang *et al*., 2015). Furthermore, the mitotic defects observed upon WAPL depletion could be rescued upon transcription inhibition, arguing that most defects associated with WAPL loss are caused by transcription-mediated events (Perea-Resa *et al*., 2020). In the present work, although transcription inhibition does not rescue the frequency of mitotic defects after depletion of Lds, it is still possible that pervasive transcription perturbs mitotic fidelity, although to a minor extent compared to defects caused by defective sister chromatid resolution. In *Drosophila* syncytial blastoderm embryos, transcription levels are low, restricted to the minor wave of zygotic gene activation (Pritchard & Schubiger, 1996). Conversely, topological issues are expected to be high due to the extremely fast genome replication. Hence, it is conceivable that in this particular system, errors in mitosis after depletion of Lds are mostly caused by decatenation defects rather than impaired transcription termination or pervasive transcription. The synergy between the emerging players in MTI as participants in other functions in chromosome architecture and/or transcriptional control, as illustrated in the present study, brings an additional challenge to the understanding of how transcription termination defects impair mitotic fidelity, and the mechanistic understanding of this interference this remains unknown.

The uncovered link between sister chromatid resolution and mitotic transcription termination reported here raises intriguing questions on how these two combined systems may have evolved and specialised across the tree of life. In mammalian cells, another SNF2 helicase-like protein, Plk1-interacting checkpoint helicase (PICH), has been documented to aid in sister chromatid resolution during anaphase by cooperating with Top2 and Top3 (Baumann *et al*, 2007; Pitchai *et al*, 2017; Spence *et al*, 2007). *Drosophila* does not have a true PICH orthologue. Lds and PICH share very low homology, except for the conserved helicase-like domains, common to the entire SNF2-like family (e.g. Lds does not have TPR domains, known to promote binding to its co-factor BEND3 (Pitchai *et al*., 2017)). Moreover, PICH is found mainly in the centromeric region (Baumann *et al*., 2007), whereas we see Lds located all over chromosomes in *Drosophila* embryos. However, it is possible that in *Drosophila*, Lds shares the function of both PICH and TTF2, the latter known to participate in mitotic transcription termination in human cells (Jiang *et al*.,2004), which have diverged in mammalian systems.

It also remains to be established how Lds can perform both functions at the mechanistic level, but it is conceivable that both mitotic transcription termination and sister chromatid resolution could rely on common mechanistic principles. In line with what has been observed for other SNF2-like proteins (Durr *et al*, 2006), Lds may induce helical torsion on DNA which could drive Pol II eviction, concomitantly favouring substrate recognition by Top2. Lds may thus be a previously unknown player in ensuring proper directionality to Top2 reactions, which is essential for efficient genome partitioning (Piskadlo & Oliveira, 2017). Further studies are required to establish how Lds acts on mitotic chromatin to impose this dual outcome.

## Materials and Methods

### Fly strains

Transgenic flies expressing EGFP-Top2 or Lds-EGFP were obtained by CRISPR-mediated mutagenesis, performed by WellGenetics Inc., using modified methods of (Kondo & Ueda, 2013). In brief, gRNA sequences GTACATCTGTTCGATGGACA[GGG] (Top2) or GGCGCCTGTAAGGACACCAT[CGG] (lds) were cloned into U6 promoter plasmid(s). Cassette EGFP containing EGFP and two homology arms were cloned into pUC57-Kan as donor template for repair. *CG10223* or *lds/ CG2684-targeting* gRNAs and hs-Cas9 were supplied in DNA plasmids, together with donor plasmid for microinjection into embryos of control strain *w[1118]*.F1 flies were screened by PCR and further validated by genomic PCR and sequencing. CRISPR generates a break upstream of *CG10223/top2 or* downstream of *lds/ CG2684* and is replaced by cassette EGFP.

All other strains used in this study were previously described and are summarised in Table 1.

**Table 1.**
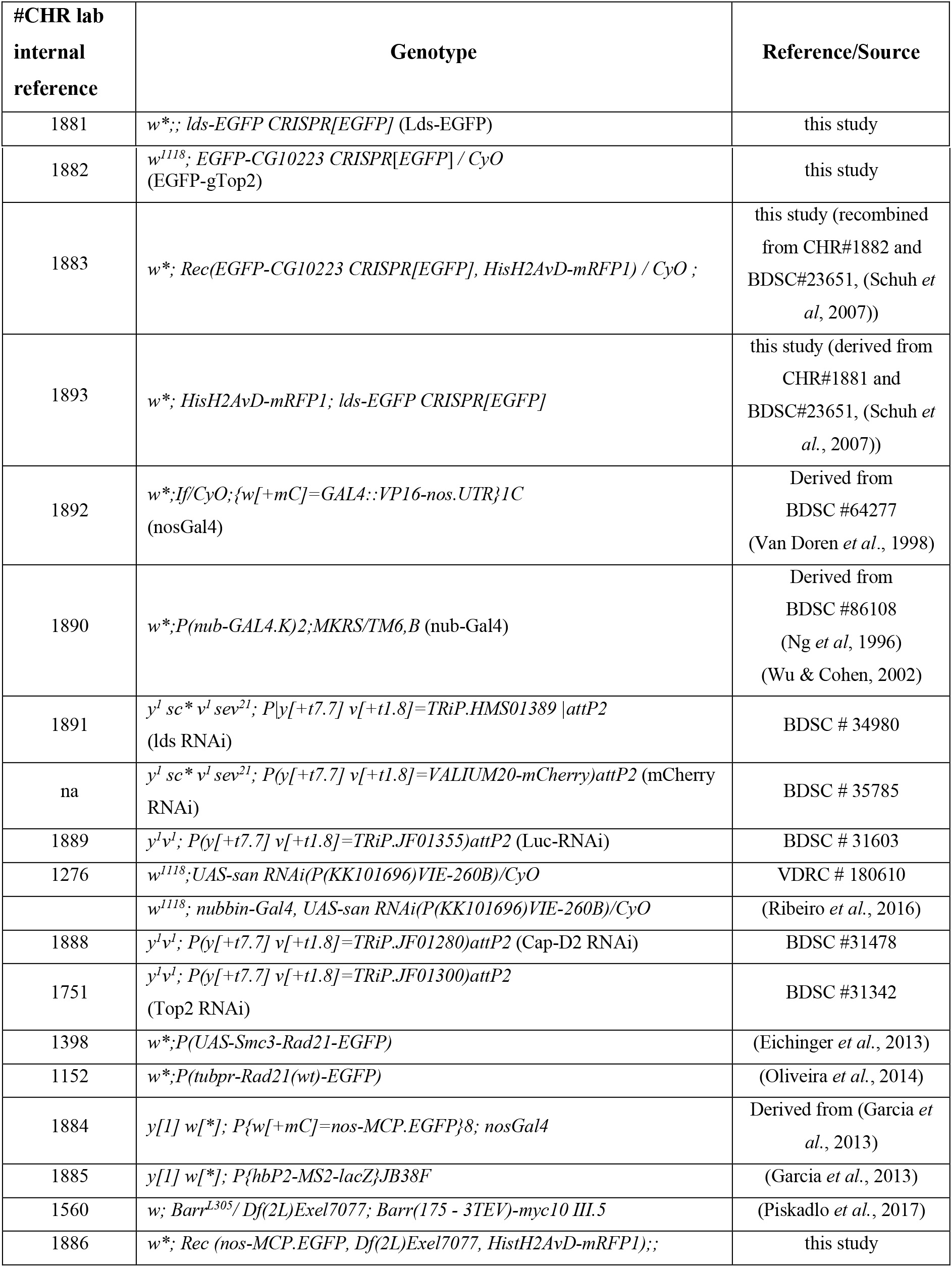
Genotype of the fly strains used in this study

### Microinjections

Embryos were prepared for imaging and microinjected as previously described (Carmo *et al*, 2019). Embryos were then microinjected with proteins/drugs at the following concentrations: alpha-amanitin (1 mg/ml), UbcH10^C114S^ (prepared as in (Oliveira *et al*., 2010), at ~30 mg/ml); ICRF-193 (280 μM); mRNA for RNAi resistant lds (1 mg/ml; prepared using mMESSAGE mMACHINE T3 transcription kit (Thermo Fisher), using Lds ORF subcloned in pRNA vector, after site-directed mutagenesis to replace the RNAi target sequence “CCGGCTCAATCTGCTAATGAA” by “TAGGCTGAACCTTCTGATGAA”).

### Microscopy

Imaginal discs and early embryos were prepared for live cell imaging as previously described (Carmo *et al*., 2019; Silva *et al*., 2018). All time-lapse movies of live embryos/wing discs (except Rad21-EGFP imaging) were obtained using Confocal Z-series stacks with a Yokogawa CSU-X Spinning Disk confocal, mounted on a Leica DMi8 microscope, with a 63x 1.3NA glycerine immersion objective, using the 488 nm and 561 nm laser lines and an Andor iXon Ultra EMCCD 1024×1024 camera. The system was controlled with Metamorph software (Molecular Devices). For imaging of Rad21-EGFP, confocal Z-series stacks were acquired on a Andor Dragonfly Spinning Disk confocal, mounted on a Nikon Ti2 microscope, with a 60x 1.2 NA water immersion objective, using the 488 nm and 561 nm laser lines and a Andor Sona sCMOS 4 MPix camera using a 1600 × 1350 FOV.

### Quantitative imaging analysis

For images analysis, z-stacks were max-intensity projected using FIJI. Mitotic errors were manually evaluated based on H2AvDmRFP1 (and Cid-EGFP in wing discs) signals. Times of anaphase and transcripts eviction were measured relative to nuclear envelope break down, defined by the time chromatin signal loses its round organisation. Transcript removal time was defined by the time MCP-EGFP labelled reporter transcripts were either no longer detected or observed outside the chromosomal region.

For quantitative analysis of chromatin bound levels of Top2/Lds (Fig 6) images were max projected, background subtracted, and Gaussian blurred (FIJI) and cropped for individual metaphases. ROIs were selected based on segmentation of the Histone mRFP1 channel, using automatic Huang threshold, to select for chromosomal area. Integrated Intensities were measured for each ROI in either Lds/Top2 and normalised to the first time point.

### Fluorescence Recovery After Photobleaching

FRAP analysis was performed in embryos from strains expressing solely the EGFP-tagged version of Lds or Top2 and HisH2AvDmRFP1 to monitor chromatin. Embryos were arrested in metaphase (UbcH10^C114S^) and subsequently microinjected with DMSO/ICRF-193. FRAP analysis was performed 10 minutes after the second microinjection to allow for stabilisation of protein accumulation on chromatin. 4 pre-bleach images were acquired every 15 sec; 14 z-stacks 0.8 um apart, followed by photobleaching of an entire metaphase with 1 pulse of 470 nm laser, using Andor’s Mosaic system (90% laser power). A total of 2-5 metaphases were bleached per embryo. Fluorescence recovery was monitored by subsequent imaging as described for pre-bleaching imaging. Quantitative analysis for fluorescence recovery for each metaphase was performed as above, measuring the integrated intensity of chromosomal regions (Hist-mRFP1-defined) over time, normalised to the image before bleach. Exponential curves were fit to a one-phase association equation (Y=Y0 + (Plateau-Y0)*(1-exp(-K*x))) using GraphPad prism 9.0 to estimate half-times of recovery (K).

### Statistical Analysis

Statistical analysis was performed using GraphPad Prism 9.0. As most data sets did not pass the normality test (D’Agostino-Pearson normality test), non-parametric tests were used including Kruskal-Wallis one-way ANOVA (for multiple comparisons) or Mann-witney. Details for each comparison can be found on the respective figure legends.

## Acknowledgements

We thank members of the Oliveira and Martinho labs for fruitful discussions. We thank the dedicated facilities at Instituto Gulbenkian de Ciência (IGC): Fly Facility, the Advanced Imaging Facility (AIF) and the Technical Support Service. Especial thanks to Stefan Heidmann (Univ. Bayreuth) and the Bloomington *Drosophila* Stock Center (BDSC) for fly stocks.

## Funding

This work was supported by the following bodies: European Research Council (ERC-2020-COG-101002391-ChromoSilence) to R.A.O., Fundação para Ciência e a Tecnologia (FCT) supporting C.C. (PD/BD/128428/2017), R.A.O. (CEECIND/01092/2017), R.G.M (PTDC/BIA-BID/1606/2020); Fundação Calouste Gulbenkian (FCG) and LISBOA-01-0145-FEDER-007654 supporting IGC’s core operation; LISBOA-01-7460145-FEDER-022170 (Congento) supporting the IGC Fly Facility; PPBI-POCI-01-0145-FEDER-022122 supporting the IGC Advanced Imaging Facility, all co-financed by FCT (Portugal) and Lisboa Regional Operational Program (Lisboa2020) under the PORTUGAL2020 Partnership Agreement (European Regional Development Fund).

## Conflict of interest

The authors declare n conflict of interest.

## Supplementary figures

**Figure S1.**
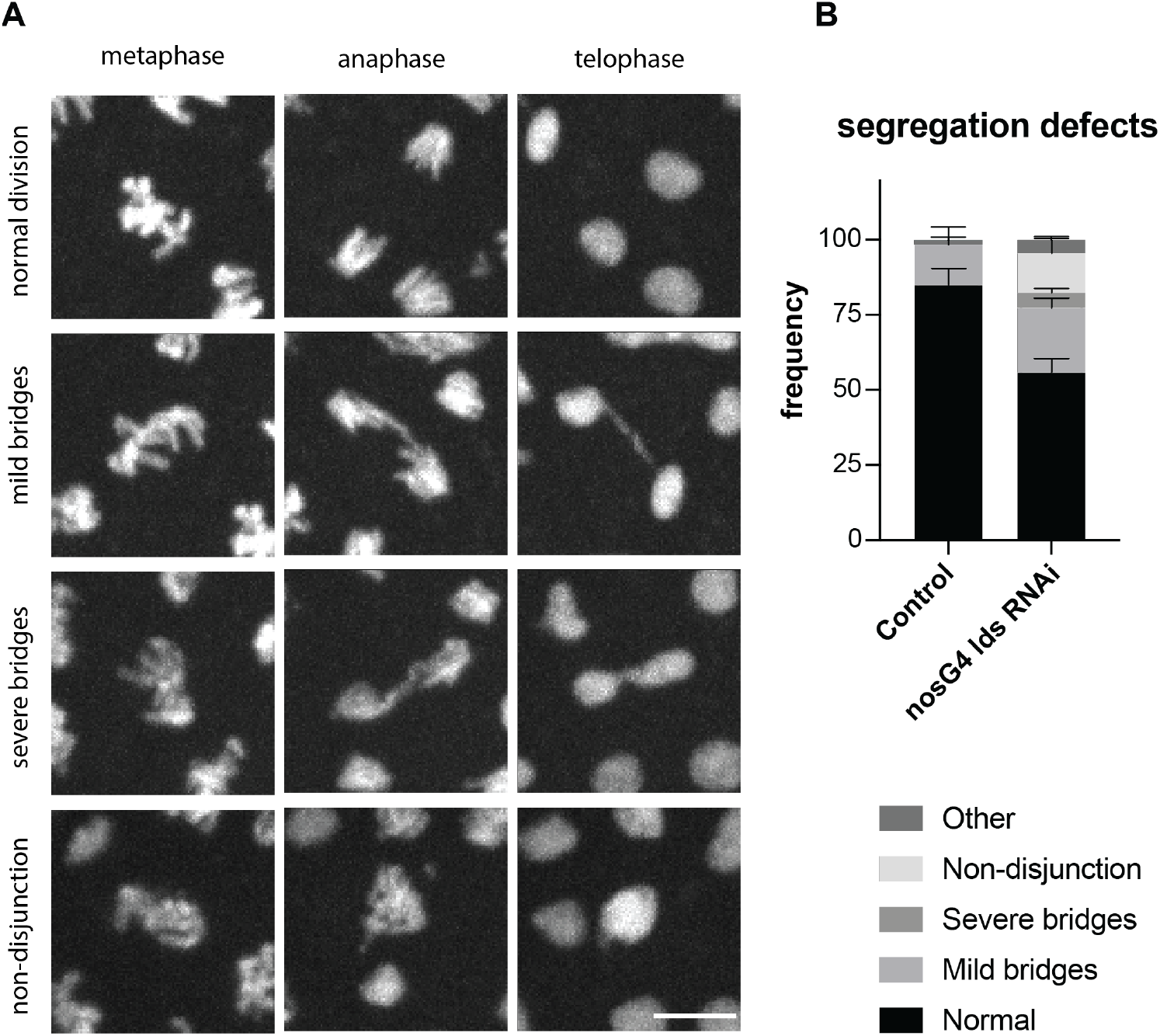
Segregation defects in *Drosophila* syncytial embryos depleted of Lds. (A) Representative images of the types of segregation errors observed upon depletion of Lds by RNAi in the germline, driven by nanos-Gal4. Scale bar is 5 μm and applies to all images. (B) Frequency of segregation defects observed in control (no RNAi, n=5 embryos) and Lds-depleted embryos (n=23 embryos). Graph represents mean and error bars SEM

**Figure S2.**
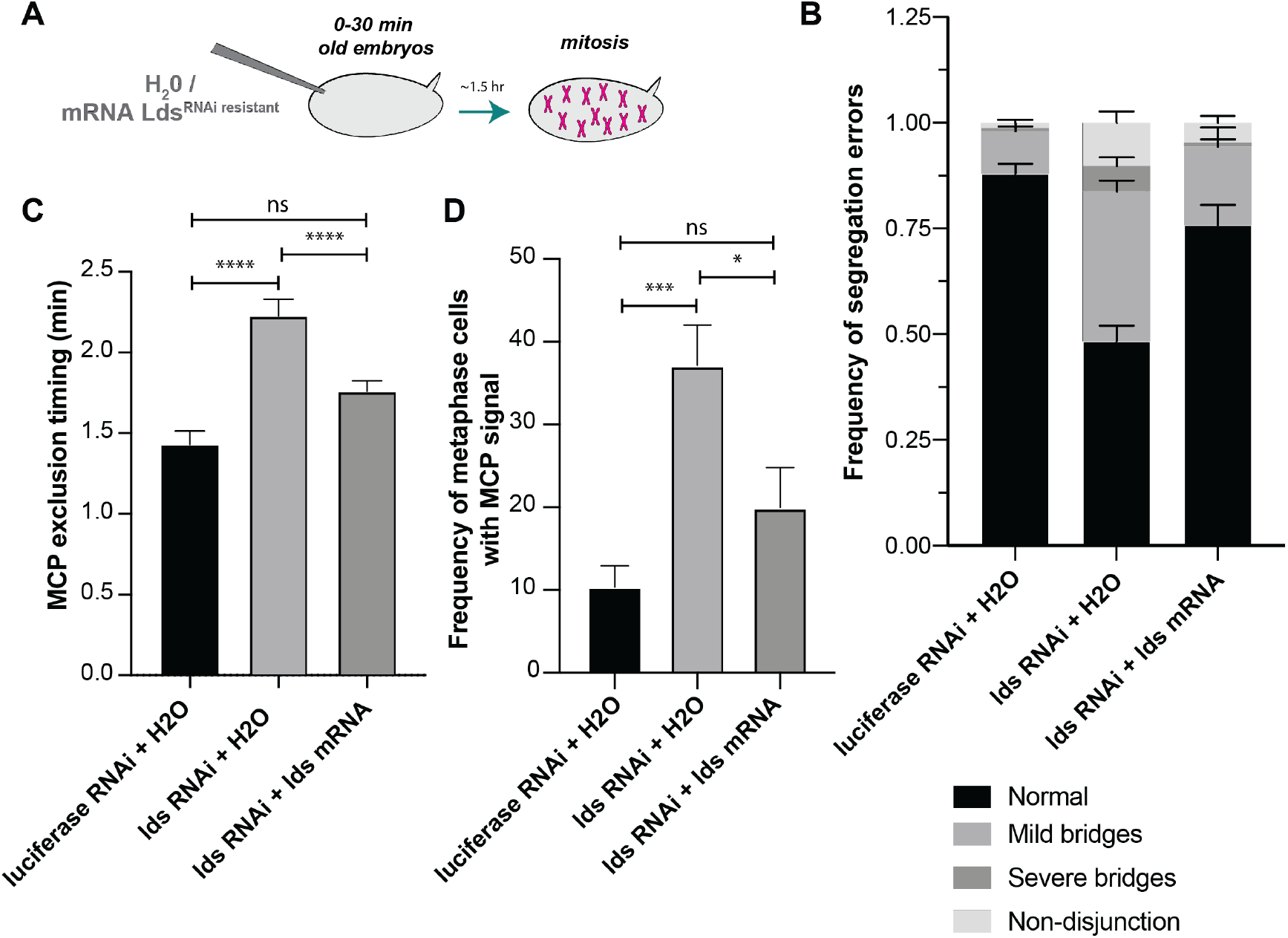
Expression of RNAi-resistant lds partially rescues the defects associated to RNAi-mediated lds depletion. **(A)** Scheme of the experimental layout for the rescue experiments: young embryos were microinjected with in vitro transcribed mRNA for Lds (RNAi resistant) and allowed to develop while translation occurs (~1.5 hr). Images were taken ~1.5 hr later to monitor transcription dynamics (using MCP-EGFP/MS2 system) and segregation defects (based on HisH2AvD-mRFP1 labelled chromatin) **(B)** Frequency of segregation defects observed in luciferase/Lds-depeted embryos, after injection of water or mRNA coding RNAi-resistant lds (luciferase-RNAi + H_2_O n=10 embryos; lds-RNAi + H_2_O n=15 embryos; lds-RNAi + mRNA lds RNAi resistant n=8 embryos) **(C)** Quantification of the timing of disappearance of transcripts from chromatin during mitosis (relative to NEBD) in the referred conditions. Bars represent mean and error bars SEM (luciferase-RNAi + H_2_O n= 171 nuclei from 7 independent embryos; lds-RNAi + H_2_O n= 191 nuclei from 9 independent embryos; lds-RNAi + mRNA lds RNAi resistant n= 360 nuclei from 17 independent embryos)

**Figure S3.**
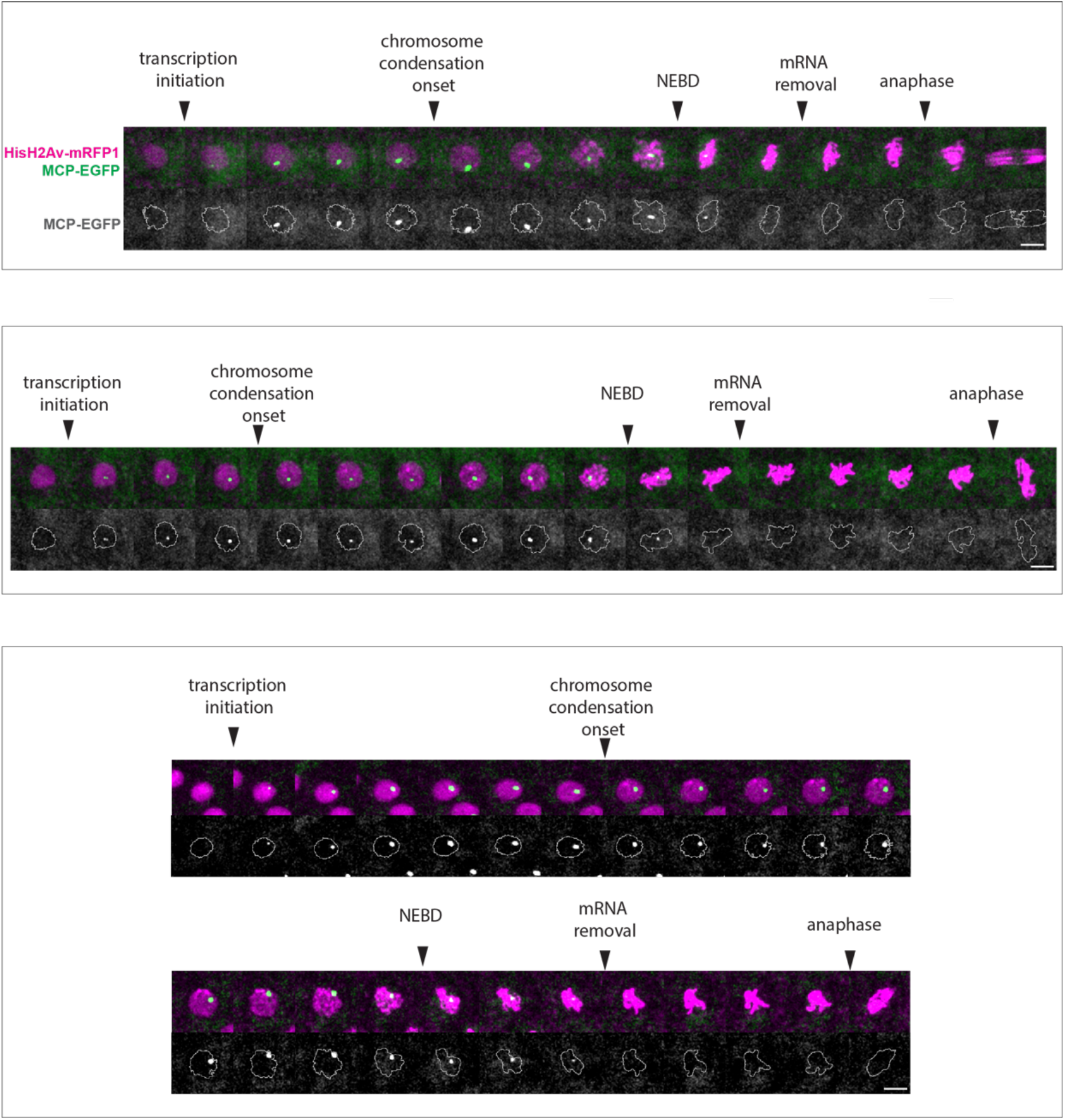
Detailed analysis of the presence of nascent transcripts on chromatin. Representative examples of nuclei undergoing mitotic divisions, with concomitant analysis of nascent transcripts (MCP-EGFP, green) and chromatin (HistH2AvDmRFP1, magenta). Bottom panels depict MCP-EGFP signal alone and chromatin area is depicted with a grey line. Note that full transcript removal is only observed after NEBD, when chromosome compaction is virtually complete. Images were taken every 30 seconds. Scale bars are 5 μm and apply to all images.

**Figure S4.**
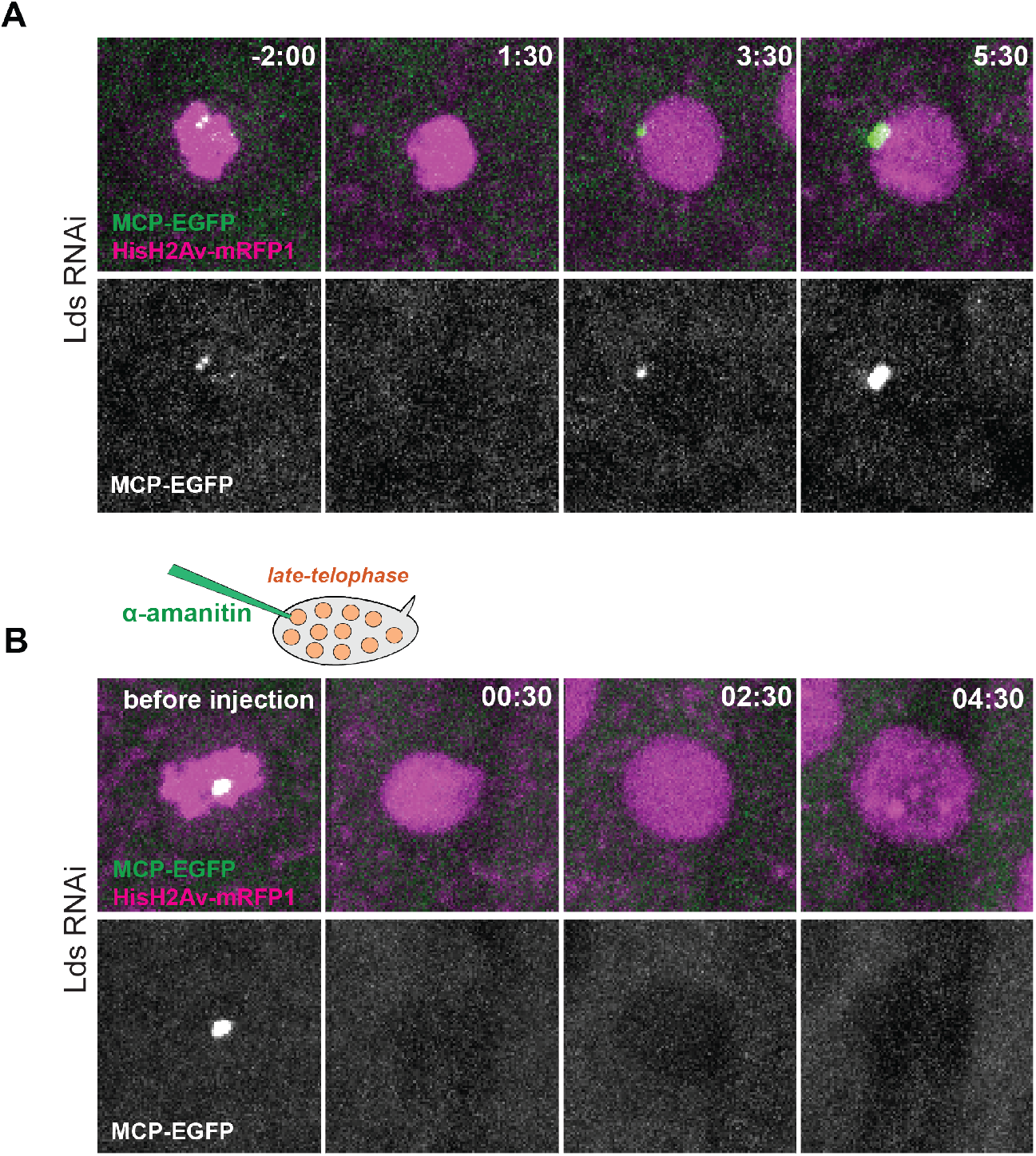
Microinjection of alpha amanitin efficiently inhibits early zygotic transcription. **(A)** Kinetics of transcription initiation in a Lds-depleted embryo (non-injected), depicting ongoing transcription in the subsequent cycle. Times are relative to the previous anaphase. DNA is labelled in magenta and nascent transcripts are in green, labelled with MCP-EGFP bound to the MS2 loops present in the transcription reporter. **(B)** Lds-depleted embryos were microinjected with 1 mg/ml alpha-amanitin in late mitosis/early interphase, and monitored for the subsequent cycle. Note that in contrast with the non-injected embryos, no transcription can be detected from the reporter gene

## References

Akoulitchev S, Reinberg D (1998) The molecular mechanism of mitotic inhibition of TFIIH is mediated by phosphorylation of CDK7. Genes Dev 12: 3541–3550

Ball CB, Parida M, Santana JF, Spector BM, Suarez GA, Price DH (2022) Nuclear export restricts Gdown1 to a mitotic function. Nucleic Acids Res 50: 1908–1926

Baumann C, Korner R, Hofmann K, Nigg EA (2007) PICH, a centromere-associated SNF2 family ATPase, is regulated by Plk1 and required for the spindle checkpoint. Cell 128: 101–114

Baxter J, Sen N, Martinez VL, De Carandini ME, Schvartzman JB, Diffley JF, Aragon L (2011) Positive supercoiling of mitotic DNA drives decatenation by topoisomerase II in eukaryotes. Science 331: 1328–1332

Bellier S, Chastant S, Adenot P, Vincent M, Renard JP, Bensaude O (1997) Nuclear translocation and carboxyl-terminal domain phosphorylation of RNA polymerase II delineate the two phases of zygotic gene activation in mammalian embryos. The EMBO journal 16: 6250–6262

Brandao HB, Paul P, van den Berg AA, Rudner DZ, Wang X, Mirny LA (2019) RNA polymerases as moving barriers to condensin loop extrusion. Proceedings of the National Academy of Sciences of the United States of America 116: 20489–20499

Carmo C, Araujo M, Oliveira RA (2019) Microinjection Techniques in Fly Embryos to Study the Function and Dynamics of SMC Complexes. Methods in molecular biology 2004: 251–268

Christensen MO, Larsen MK, Barthelmes HU, Hock R, Andersen CL, Kjeldsen E, Knudsen BR, Westergaard O, Boege F, Mielke C (2002) Dynamics of human DNA topoisomerases IIalpha and IIbeta in living cells. The Journal of cell biology 157: 31–44

Davidson IF, Bauer B, Goetz D, Tang W, Wutz G, Peters JM (2019) DNA loop extrusion by human cohesin. Science 366: 1338–1345

Durr H, Flaus A, Owen-Hughes T, Hopfner KP (2006) Snf2 family ATPases and DExx box helicases: differences and unifying concepts from high-resolution crystal structures. Nucleic Acids Res 34: 4160–4167

Eichinger CS, Kurze A, Oliveira RA, Nasmyth K (2013) Disengaging the Smc3/kleisin interface releases cohesin from Drosophila chromosomes during interphase and mitosis. The EMBO journal 32: 656–665

Farcas AM, Uluocak P, Helmhart W, Nasmyth K (2011) Cohesin’s concatenation of sister DNAs maintains their intertwining. Mol Cell 44: 97–107

Gandhi R, Gillespie PJ, Hirano T (2006) Human Wapl is a cohesin-binding protein that promotes sister-chromatid resolution in mitotic prophase. Current biology: CB 16: 2406–2417

Ganji M, Shaltiel IA, Bisht S, Kim E, Kalichava A, Haering CH, Dekker C (2018) Real-time imaging of DNA loop extrusion by condensin. Science 360: 102–105

Garcia HG, Tikhonov M, Lin A, Gregor T (2013) Quantitative imaging of transcription in living Drosophila embryos links polymerase activity to patterning. Current biology: CB 23: 2140–2145

Gebara MM, Sayre MH, Corden JL (1997) Phosphorylation of the carboxy-terminal repeat domain in RNA polymerase II by cyclin-dependent kinases is sufficient to inhibit transcription. J Cell Biochem 64: 390–402

Gimenez-Abian JF, Sumara I, Hirota T, Hauf S, Gerlich D, de la Torre C, Ellenberg J, Peters JM (2004) Regulation of sister chromatid cohesion between chromosome arms. Current biology: CB 14: 1187–1193

Girdham CH, Glover DM (1991) Chromosome tangling and breakage at anaphase result from mutations in lodestar, a Drosophila gene encoding a putative nucleoside triphosphate-binding protein. Genes Dev 5: 1786–1799

Gottesfeld JM, Forbes DJ (1997) Mitotic repression of the transcriptional machinery. Trends in biochemical sciences 22: 197–202

Hou F, Chu CW, Kong X, Yokomori K, Zou H (2007) The acetyltransferase activity of San stabilizes the mitotic cohesin at the centromeres in a shugoshin-independent manner. The Journal of cell biology 177: 587–597

Jiang Y, Liu M, Spencer CA, Price DH (2004) Involvement of transcription termination factor 2 in mitotic repression of transcription elongation. Mol Cell 14: 375–385

Kim Y, Shi Z, Zhang H, Finkelstein IJ, Yu H (2019) Human cohesin compacts DNA by loop extrusion. Science 366: 1345–1349

Kleinschnitz K, Viessmann N, Jordan M, Heidmann SK (2020) Condensin I is required for faithful meiosis in Drosophila males. Chromosoma 129: 141–160

Kondo S, Ueda R (2013) Highly improved gene targeting by germline-specific Cas9 expression in Drosophila. Genetics 195: 715–721

Kueng S, Hegemann B, Peters BH, Lipp JJ, Schleiffer A, Mechtler K, Peters JM (2006) Wapl controls the dynamic association of cohesin with chromatin. Cell 127: 955–967

Liang HL, Nien CY, Liu HY, Metzstein MM, Kirov N, Rushlow C (2008) The zinc-finger protein Zelda is a key activator of the early zygotic genome in Drosophila. Nature 456: 400–403

Liang K, Woodfin AR, Slaughter BD, Unruh JR, Box AC, Rickels RA, Gao X, Haug JS, Jaspersen SL, Shilatifard A (2015) Mitotic Transcriptional Activation: Clearance of Actively Engaged Pol II via Transcriptional Elongation Control in Mitosis. Mol Cell 60: 435–445

Liu M, Xie Z, Price DH (1998) A human RNA polymerase II transcription termination factor is a SWI2/SNF2 family member. J Biol Chem 273: 25541–25544

Long JJ, Leresche A, Kriwacki RW, Gottesfeld JM (1998) Repression of TFIIH transcriptional activity and TFIIH-associated cdk7 kinase activity at mitosis. Mol Cell Biol 18: 1467–1476

Losada A, Hirano M, Hirano T (1998) Identification of Xenopus SMC protein complexes required for sister chromatid cohesion. Genes Dev 12: 1986–1997

Nakazawa N, Arakawa O, Yanagida M (2019) Condensin locates at transcriptional termination sites in mitosis, possibly releasing mitotic transcripts. Open biology 9: 190125

Ng M, Diaz-Benjumea FJ, Vincent JP, Wu J, Cohen SM (1996) Specification of the wing by localized expression of wingless protein. Nature 381: 316–318

Oliveira RA, Hamilton RS, Pauli A, Davis I, Nasmyth K (2010) Cohesin cleavage and Cdk inhibition trigger formation of daughter nuclei. Nature cell biology 12: 185–192

Oliveira RA, Heidmann S, Sunkel CE (2007) Condensin I binds chromatin early in prophase and displays a highly dynamic association with Drosophila mitotic chromosomes. Chromosoma 116: 259–274

Oliveira RA, Kotadia S, Tavares A, Mirkovic M, Bowlin K, Eichinger CS, Nasmyth K, Sullivan W (2014) Centromere-independent accumulation of cohesin at ectopic heterochromatin sites induces chromosome stretching during anaphase. PLoS biology 12: e1001962

Perea-Resa C, Bury L, Cheeseman IM, Blower MD (2020) Cohesin Removal Reprograms Gene Expression upon Mitotic Entry. Mol Cell 78: 127–140 e127

Piskadlo E, Oliveira RA (2016) Novel insights into mitotic chromosome condensation. F1000Res 5

Piskadlo E, Oliveira RA (2017) A Topology-Centric View on Mitotic Chromosome Architecture. Int J Mol Sci 18

Piskadlo E, Tavares A, Oliveira RA (2017) Metaphase chromosome structure is dynamically maintained by condensin I-directed DNA (de)catenation. Elife 6

Pitchai GP, Kaulich M, Bizard AH, Mesa P, Yao Q, Sarlos K, Streicher WW, Nigg EA, Montoya G, Hickson ID (2017) A novel TPR-BEN domain interaction mediates PICH-BEND3 association. Nucleic Acids Res 45: 11413–11424

Pommier Y, Sun Y, Huang SN, Nitiss JL (2016) Roles of eukaryotic topoisomerases in transcription, replication and genomic stability. Nature reviews Molecular cell biology 17: 703–721

Price DH, Sluder AE, Greenleaf AL (1987) Fractionation of transcription factors for RNA polymerase II from Drosophila Kc cell nuclear extracts. J Biol Chem 262: 3244–3255

Pritchard DK, Schubiger G (1996) Activation of transcription in Drosophila embryos is a gradual process mediated by the nucleocytoplasmic ratio. Genes Dev 10: 1131–1142

Ribeiro AL, Silva RD, Foyn H, Tiago MN, Rathore OS, Arnesen T, Martinho RG (2016) Naa50/San-dependent N-terminal acetylation of Scc1 is potentially important for sister chromatid cohesion. Sci Rep 6: 39118

Rong Z, Ouyang Z, Magin RS, Marmorstein R, Yu H (2016) Opposing Functions of the N-terminal Acetyltransferases Naa50 and NatA in Sister-chromatid Cohesion. J Biol Chem 291: 19079–19091

Rothe M, Pehl M, Taubert H, Jackle H (1992) Loss of gene function through rapid mitotic cycles in the Drosophila embryo. Nature 359: 156–159

Ryoo HD, Gorenc T, Steller H (2004) Apoptotic cells can induce compensatory cell proliferation through the JNK and the Wingless signaling pathways. Developmental cell 7: 491–501

Schuh M, Lehner CF, Heidmann S (2007) Incorporation of Drosophila CID/CENP-A and CENP-C into Centromeres during Early Embryonic Anaphase. Current biology: CB 17: 237–243

Segil N, Guermah M, Hoffmann A, Roeder RG, Heintz N (1996) Mitotic regulation of TFIID: inhibition of activator-dependent transcription and changes in subcellular localization. Genes Dev 10: 2389–2400

Sen N, Leonard J, Torres R, Garcia-Luis J, Palou-Marin G, Aragon L (2016) Physical Proximity of Sister Chromatids Promotes Top2-Dependent Intertwining. Mol Cell 64: 134–147

Shermoen AW, O’Farrell PH (1991) Progression of the cell cycle through mitosis leads to abortion of nascent transcripts. Cell 67: 303–310

Silva RD, Mirkovic M, Guilgur LG, Rathore OS, Martinho RG, Oliveira RA (2018) Absence of the Spindle Assembly Checkpoint Restores Mitotic Fidelity upon Loss of Sister Chromatid Cohesion. Current biology: CB 28: 2837–2844 e2833

Spence JM, Phua HH, Mills W, Carpenter AJ, Porter AC, Farr CJ (2007) Depletion of topoisomerase IIalpha leads to shortening of the metaphase interkinetochore distance and abnormal persistence of PICH-coated anaphase threads. Journal of cell science 120: 3952–3964

Staudt N, Fellert S, Chung HR, Jackle H, Vorbruggen G (2006) Mutations of the Drosophila zinc finger-encoding gene vielfaltig impair mitotic cell divisions and cause improper chromosome segregation. Mol Biol Cell 17: 2356–2365

Sumara I, Vorlaufer E, Gieffers C, Peters BH, Peters JM (2000) Characterization of vertebrate cohesin complexes and their regulation in prophase. The Journal of cell biology 151: 749–762

Szalontai T, Gaspar I, Belecz I, Kerekes I, Erdelyi M, Boros I, Szabad J (2009) HorkaD, a chromosome instability-causing mutation in Drosophila, is a dominant-negative allele of Lodestar. Genetics 181: 367–377

Tadros W, Lipshitz HD (2009) The maternal-to-zygotic transition: a play in two acts. Development 136: 3033–3042

Tavormina PA, Come MG, Hudson JR, Mo YY, Beck WT, Gorbsky GJ (2002) Rapid exchange of mammalian topoisomerase II alpha at kinetochores and chromosome arms in mitosis. The Journal of cell biology 158: 23–29

Wiegard A, Kuzin V, Cameron DP, Grosser J, Ceribelli M, Mehmood R, Ballarino R, Valant F, Grochowski R, Karabogdan I et al (2021) Topoisomerase 1 activity during mitotic transcription favors the transition from mitosis to G1. Mol Cell 81: 5007–5024 e5009

Williams BC, Garrett-Engele CM, Li Z, Williams EV, Rosenman ED, Goldberg ML (2003) Two putative acetyltransferases, san and deco, are required for establishing sister chromatid cohesion in Drosophila. Current biology: CB 13: 2025–2036

Wu J, Cohen SM (2002) Repression of Teashirt marks the initiation of wing development. Development 129: 2411–2418

Xie Z, Price D (1997) Drosophila factor 2, an RNA polymerase II transcript release factor, has DNA-dependent ATPase activity. J Biol Chem 272: 31902–31907

Xie Z, Price DH (1996) Purification of an RNA polymerase II transcript release factor from Drosophila. J Biol Chem 271: 11043–11046

Xie Z, Price DH (1998) Unusual nucleic acid binding properties of factor 2, an RNA polymerase II transcript release factor. J Biol Chem 273: 3771–3777

Xing H, Vanderford NL, Sarge KD (2008) The TBP-PP2A mitotic complex bookmarks genes by preventing condensin action. Nature cell biology 10: 1318–1323

